# 10 recommendations for strengthening citizen science for improved societal and ecological outcomes: A co-produced analysis of challenges and opportunities in the 21^st^ century

**DOI:** 10.1101/2025.08.13.670232

**Authors:** Jack Nunn, Håkon da Silva Hyldmo, Lauren McKnight, Heather McCulloch, Jennifer Lavers, Julie Old, Laura Smith, Nicola Grobler, Cheryl Tan Kay Yin, Wing Yan Chan, Candice Raeburn, Nittya S. M. Simard, Adam Kingsley Smith, Sam Van Holsbeeck, Eleanor Drinkwater, Kit Prendergast, Emma Burrows, Christopher L. Lawson

## Abstract

Citizen science plays an increasingly important role in generating scientific knowledge and supporting environmental and social action. However, its potential to address complex global challenges remains underutilised. This study explores how to improve citizen science by involving the public in all stages of scientific research. Using participatory research methods, we conducted online surveys and group discussions with researchers, citizen scientists, and Indigenous people. Thematic coding was used to identify key challenges, opportunities, and best practices to enhance citizen science initiatives. Additionally, nine case studies were reported using the Standardised Data on Initiatives (STARDIT) reporting tool.

The study identifies key strategies for enhancing participant engagement and retention in citizen science initiatives. Findings underscore the importance of inclusive, evidence-informed approaches such as targeted outreach, fair compensation, tailored support, and co-creation practices. Ensuring data quality and fostering trust require adherence to FAIR data principles, transparent validation and sharing processes, and the establishment of ethical research partnerships. Persistent challenges include short-term funding, which undermines long-term project sustainability, and the lack of centralized support for ethics and project management. Formal recognition of citizen scientists through co-authorship, standardized training, and professional development opportunities can further strengthen involvement and build capacity. Finally, emerging technologies—including AI and open data platforms—present opportunities to scale and improve efficiency, provided they are implemented with appropriate ethical safeguards and investment.

Drawing together these insights, we provide 10 actionable recommendations for citizen science in the 21^st^ century. These highlight the importance of embedding citizen science in national research infrastructure, education, and policy, alongside consistent evaluation and reporting, to improve its inclusivity, longevity, and impact. We conclude by arguing that as the world confronts climate change, public health crises, and biodiversity loss, broader public involvement in science is key for equitable, efficient and evidence-informed responses.

## Introduction

Throughout human history, knowledge-sharing has been a participatory process. The Universal Declaration of Human Rights recognises this by stating that everyone has the right to “receive and impart information and ideas through any media and regardless of frontiers” (1). However, in the past century there has been a real or perceived division between those in academia who are paid to ‘do science’, and those who are outside of academia or unpaid, including citizen scientists (2–4). Citizen science is a ‘flexible concept’, but has clear principles and values associated with it. Acknowledging that the concept of citizen science and its terminology are complex and flexible, we here define ‘citizen science’ to mean ‘the active involvement of the public at any stage of scientific research, with the aim to increase scientific knowledge, including by using the scientific method’ (5).

Citizen science often emphasises partnerships among the public, non-professional scientists and professional scientists, with the aim of democratising the scientific process by making it more inclusive, accessible, and impactful. However, the concept of citizen science is not restricted to questions about who is ‘a professional’, who is being paid and who isn’t, nor is it limited to the data collection stage of research. Citizen science as a concept is an open approach to science, which enables people to be involved in all stages of the scientific method, including identifying gaps in our knowledge and research priorities, developing research questions, designing methods, gathering and analysing data, communicating results, and implementing learnings. Further, the goals of citizen science can vary, for example to collect more scientific data or to improve public awareness of scientific issues (6).

Citizen science and its associated methods are not limited to certain disciplines. Disciplines such as health, environmental and educational sciences all use varied language to describe the same concepts of sharing power and responsibility with the public, and the democratisation of all stages of research (7). While concepts of public involvement in health research and healthcare have been well-defined for a number of decades (8), more precise definitions and methodologies are still being discussed and refined across diverse disciplines (9,10). This includes health research (11) and the ‘one health’ approach (12); environmental research (13,14); evidence synthesis (15,16); urban planning (17); education (18); law (19); policy (20,21); and economics and budgeting (22). Across all these disciplines, we can distinguish between scientific knowledge (what we know), the scientific method (how we try to answer questions), and the specific tools used to conduct the scientific method. Many different tools within citizen science can be applied across disciplines.

Citizen science is connected to many other related concepts (10,23). A core concept of the many manifestations of citizen science is ‘critical pedagogy’, which was defined by Paulo Freire as seeing “the world not as a static reality, but as a reality in process, in transformation” (24). This inspired the development of ‘participatory action research’ as an approach to research methods where “communities of inquiry and action evolve and address questions and issues that are significant for those who participate as co-researchers” (25). The philosophy of the ‘open knowledge’ movement and the concept of ‘open science’ are also intrinsically connected to citizen science (26). In 2021, the United Nations Educational, Scientific and Cultural Organization (UNESCO) published the ‘Recommendation on Open Science’, which recognised that “science is a global public good that belongs to all of humanity” and is intended to open “the processes of scientific knowledge creation, evaluation and communication to societal actors beyond the traditional scientific community” (27). The intention of the Recommendation is that by promoting open science, science becomes “more accessible, inclusive and transparent” and “furthers the right of everyone to share in scientific advancement and its benefits” (28). Further, the World Health Organisation identifies the need for “a meaningful whole-of-society approach and social participation” in universal health coverage, and health and well-being (29).

### Motivation for this discussion paper

Citizen science can improve how we conduct research and involve people in helping understand our universe. Effective and inclusive citizen science requires partnerships, resourcing and support from multiple organisations. The benefits of citizen science can also include improved public involvement and scientific literacy, improved large-scale data collection for biodiversity monitoring and early threat detection, fostering of environmental stewardship, collaboration between professional scientists and the public, empowering communities at greater risk of exploitation, inspiring people to study and work in science, and building public trust in science through transparency and inclusivity (30–34). Previous research has demonstrated the importance of creating enabling conditions for inclusive citizen science by building people’s capacity through training and development, developing institutional partnerships, providing infrastructure, long-term funding, and accessible technologies that facilitate data collection and analysis (10,35,36).

However, initiating and leading citizen science projects presents challenges such as ensuring data quality, retaining participant who are involved, and addressing ethical concerns like consent and data ownership (37). Additionally, those leading them face difficulties related to inclusivity, communication, funding, and balancing scientific control with meaningful public involvement (38–40). The challenges faced by those leading citizen science projects and those involved in them led to the commissioning of this discussion paper that sought to involve both researchers and other people involved in citizen science. This discussion paper aims to identify key themes, challenges and solutions for advancing and improving citizen science for all involved. A participatory action research approach was adopted when scoping and writing this article, which allowed the co-authors to shape the discussion to focus on areas they identified as priorities. We present our findings in this discussion paper, summarise what was learned, and make recommendations based on the data collected.

This research was guided by participatory action research paradigms, which guide initiatives by aiming to involve multiple stakeholders in every aspect of their development and evaluation (41–43). It was co-created with citizen science experts from health research, environmental research, and education across various countries. The aim was to identify diverse expert advice, best practices, and lessons learned for strengthening the societal and environmental contributions of citizen science. Citizen science experts and practitioners collaboratively designed a survey, analysed the results, and co-authored key recommendations based on the participants’ collective knowledge. Given the participatory co-designed protocol used in this study, we have included explanations in the methods for which tasks were completed by different co-authors.

## Methods

### Overview

The data collection was divided into two distinct rounds (Fig 1). Round One of the survey was co-designed by project coordinators and included an open invitation for co-authorship in the survey (following guidelines of the Committee of Publication Ethics (COPE)(44)). Contributions by co-authors were transparently reported using Standardised Data on Initiatives (STARDIT), which is an open-access data-sharing system for standardising how information about initiatives is reported (45). Co-authors were invited to comment on the data, analysis methods, and to identify any perceived gaps in the data collected.

**Fig 1:** Schematic overview of co-production process. Overview of the co-production process used in the production of this paper.

In Round Two, co-authors refined the research scope and survey questions, expanded data categories, and contributed to research planning. Communication modes included shared collaborative documents, video calls, and text-based discussion platforms. Tasks included data analysis, text searches, mixed-methods design, advising on ethics, and the creation of STARDIT reports about the case studies. They also mapped themes, identified case studies and completed a cross-case analysis. Co-authors collaborated on writing and reviewing the discussion paper by working on shared drafts and providing feedback to ensure a rigorous and inclusive research process.

The co-creation process for this article invited people to volunteer their time, with the incentive of being a co-author on a peer-reviewed publication. However, co-authors agreed during the co-creation process that some people should be paid for some tasks. Further information about this is provided in the Supplementary Information and the STARDIT report about this article (46).

### Data collection

#### Online surveys

Before Round One data was collected, the purpose and content of the questions were discussed and refined with the lead author (JN) and members of the Australian Academy of Sciences Australia. A structured questionnaire was created using an online survey platform (Google Forms), which could be accessed and completed by anyone in the world. It collected demographic data and featured open-ended questions about definitions of citizen science; areas of research or practice where citizen science could be better harnessed; examples of best practice (including an option to share URLs); challenges, opportunities and barriers faced by citizen science projects; and areas for growth and improvement (Appendix 1). The survey also asked respondents if they would like to be contacted about being co-authors.

The Round One survey was shared on 22 July 2024, and data was collected for 67 days. The Australian Academy of Sciences Australia and Science for All shared the survey link via email newsletters and on multiple social media platforms, including Facebook, X (formerly Twitter), and LinkedIn. Co-authors identified in the Round One survey were then invited to an online ‘kick-off’ meeting. Several follow-up meetings and shared documents were used to analyse the responses from Round One, and to improve and refine data collection for Round Two. The wording of some questions was changed, and further questions were added to Round Two, which was shared using the same methods as Round One on 10 October 2024 for 21 days (Appendix 2). A final survey about people’s experiences of being involved in the project and using STARDIT was collected from 7 – 30 July 2025 (Appendix 3). All participants provided written informed consent to participate in the study. In addition to the survey data collected, 20 online meetings were held. Additional ideas that the group felt were important to the wider discussion were also added after the final data collection.

### Data analysis

#### Qualitative analysis

The qualitative analysis was conducted in six stages.

##### Stage 1: Dataset preparation

Personally identifiable information, including names, contact details, and workplace references was removed. Each question was given an ID, and each survey respondent was assigned a unique identifier to allow re-identification if necessary. A master redacted dataset was created and securely shared with co-authors.

##### Stage 2: Inductive code creation

Based on survey responses, related terms of an issue, idea or topic were identified as themes for categorising responses, and a draft code set was developed. ‘Coding’ here refers to identifying and labelling key ideas or themes emerging from the data in a systematic way. This draft code was tested against survey responses, and an “Other – self-describe” category was added for flexibility. The coding system was refined through an iterative process that included co-author review, which ensured completeness and usability for survey responses. The final draft of the codes was shared for analysis to categorise each response as either a ‘challenge’ or ‘opportunity’ within one or more themes.

##### Stage 3: Codifying responses

Survey responses were randomly assigned to individuals for coding. Each team member applied relevant themes and sub-themes to their allocated data subset. Potential conflicts of interest were identified and addressed by re-allocating response data. A live-shared analysis template was used (Google Docs), and missing themes were added under the “Other” category as needed.

##### Stage 4: Collating and refining codified responses

All coded responses were consolidated into a single document. The code set was refined as new themes and sub-themes emerged. Any newly identified themes from the “Other” category were incorporated into the dataset. A shared document was maintained for collaboration.

##### Stage 5: Final coding and quality assurance

A final round of coding was conducted using the updated code set. Co-authors peer reviewed the code set by validating a subset of coded responses. Discrepancies were discussed and resolved. The lead author conducted a final quality check on 20% of responses, and any major inconsistencies were recoded by another author. Group discussions during video calls and text-based discussions also helped resolve coding decisions.

##### Stage 6: Preparation for write-up

The coded themes were converted into prose for the Supporting Information and structured into readable sections for the main article. The Ten Principles of Citizen Science as described by the European Citizen Science Association (47) and the Australian Citizen Science Association (5) were then mapped to the relevant themes. Co-authors developed 10 recommendations for action in citizen science programs that directly respond to the main emergent themes in the data.

#### Quantitative analysis

Quantitative analysis was used to identify and count keywords in each theme that were identified by the qualitative analysis. We analysed responses from the two rounds of surveys and analysed various aspects of citizen science practices were examined. For open-ended questions, text analysis was performed using Python’s Natural Language Toolkit (NLTK). Responses underwent preprocessing, including lowercase conversion, special character removal, and tokenisation. Theme modelling was implemented using Latent Dirichlet Allocation (LDA) and SciKit-Learn’s LatentDirichletAllocation and CountVectorizer for text vectorization. Custom stop words were employed and domain-specific stop words removed to improve theme coherence and analysis quality. Word frequency analysis was conducted using NLTK’s FreqDist. Data visualisation was performed using matplotlib and seaborn libraries, which generated response rate charts and word frequency distributions.

For Round 2, additional task-specific analyses were visualised using individual bar charts. The analysis was conducted using Python 3.13 with key libraries including NLTK, scikit-learn, pandas, matplotlib, and seaborn. All analyses were performed in a Jupyter notebook environment to ensure reproducibility and documentation. The combination of these methods provided a comprehensive understanding of patterns and trends in participant responses across both survey rounds.

For the final survey on people’s experience of being involved in this paper, we calculated the percentage of each response option in each question.

### Case studies and data reporting

Survey respondents provided examples of citizen science initiatives that exhibited best practices to benefit society. The co-authors conducted desktop research and online searches to gather supplementary publicly accessible information and data on these best-practice examples.

After analysis of the emerging themes from the qualitative data, the co-authors collaboratively selected citizen science initiatives that captured the breadth and variety in the material or were illustrative of certain themes. We have included these illustrative case studies to demonstrate with real-world examples some of the themes we identified from the survey data (26). The initiatives were analysed in more detail by extracting and structuring their data using STARDIT data fields (45) (including aims, outcomes, data sharing and budgets), leading to the creation of STARDIT reports for each case study.

This entire process was conducted collaboratively and transparently, with co-authors verifying data accuracy. Organisations and individuals directly involved in each initiative were contacted via email and given the opportunity to review, amend, or improve their respective STARDIT reports.

## Results

### Demographic data

A total of 46 respondents participated in the survey (28 in Round One, 18 in Round Two) and shared a total of 16,247 words in open text fields, with an average of approximately 350 words per respondent. Most respondents lived in Australia (80%), two (4%) of which were Aboriginal or Torres Strait Islander persons. Most respondents considered themselves to be ‘academic’ and ‘have experience with academic writing’ (76%), including professional researchers paid to do research (65%). Most also reported that they had experience as a volunteer researcher contributing to citizen science (65%) (Table S1). All 46 respondents provided an email address and 34 answered that they wanted to contribute as a co-author. A total of 29 people completed the final feedback on the experience of being involved. A total of 18 respondents co-authored the final version of this paper (in line with the Committee on Publication Ethics Authorship guidelines (44)), and 6 asked to be acknowledged for their support. In total, people estimated that they volunteered 1385 hours on this project.

### Data on payment for expertise in citizen science projects

Round Two gathered data on tasks in citizen science projects, and 17 out of 18 respondents provided answers about whether they have completed paid or unpaid work in citizen science, as well as which tasks they think should be paid (Fig 2a). Most participants had experience contributing as a co-author, dissemination including conference presentations, and funding applications, and these tasks were predominantly performed without payment (Fig 2a). The remaining seven tasks were more commonly paid for, with the highest proportion of paid positions observed in project management, planning and design (Fig 2a).

**Fig 2:** Results from Round Two survey on respondents’ experience and opinions regarding paid tasks in citizen science projects. **(a)** Tasks previously performed by respondents in citizen science projects, indicating whether they were undertaken as paid or unpaid work (multiple responses allowed per task). **(b)** Respondents’ opinions on which tasks in citizen science projects should be financially compensated, reflecting perceptions of task value, effort, or required expertise.

When asked about which tasks should be paid for, respondents showed the strongest support for data analysis, project management, and ethics applications (Fig 2b). While data collection was one of the most commonly performed paid and unpaid activities, there was relatively less support for making this a paid task (Fig 2b).

### Main themes from data

Eleven main themes with at least 15 responses per theme were identified through manual coding. The themes and a summary of how they relate to challenges and opportunities for citizen science is presented here.

- **Recruitment and Awareness (43 responses)**: Challenges include difficulties in recruiting people to be involved, a lack of public awareness about citizen science projects, structural barriers (such as no payment for those involved), and lack of support for researchers.
- **Data Collection and Confidence (38 responses)**: Concerns about real or perceived poor data quality are widespread. Poor data continuity is also a significant challenge due to issues with retention, funding, and establishing long-term expert collaboration.
- **Project Support and Capacity (26 responses)**: There is a lack of centralised support for operationalising robust citizen science projects, such as project management, ethics, and usable online infrastructure for citizen scientists.
- **Attitudes (25 responses)**: Citizen science is hindered by the perception of science as ‘boring’ and unreliable. Universities may show disinterest due to a lack of direct funding for citizen science, and scientists or medical practitioners may view citizen science as merely ‘outreach’ or ‘engagement’ rather than an approach to research.
- **Involvement and Retention (24 responses)**: Challenges include retaining existing citizen scientists, delays between data collection and research outcomes, lack of funding and resources for ongoing involvement, and a reduction in data quality due to the loss of citizen science participants.
- **Individual Capability and Training (23 responses)**: Professional researchers often lack the training to involve people, identify appropriate collaborating organisations, and design projects. Providing adequate training for people involved in citizen science is also a challenge.
- **Funding (23 responses)**: There is a general lack of direct funding for citizen science as a methodology across all fields. Short-term grants limit research partnerships and negatively affect data continuity and project longevity.
- **Inclusion and Access (20 responses)**: People involved in citizen science can be seen as merely data collectors without authentic involvement in the whole research process. Ethical data sharing and intellectual property rights pose ongoing challenges, especially when working with Indigenous participants.
- **Recognition of Volunteers (20 responses)**: People involved in citizen science are often under-acknowledged and require additional support or recognition to ensure they feel that their contributions are valued. Participants may lack access to the data they collected or contributed to.
- **Technology and Innovation (16 responses)**: There are too many redundant smartphone applications, and novel applications are technically difficult and expensive to build and maintain. Participants can also find it difficult to identify relevant citizen science smartphone applications. Artificial Intelligence can increase the scope of citizen science.
- **Educational Institutions (15 responses)**: There is a lack of awareness of the benefits of citizen science for both students and staff, which combined with few other incentives, means it is difficult to maintain continued student involvement in projects.

There were many overlaps and interplays of challenges and opportunities amongst themes. For example, attitudes towards citizen science emerged as its own theme, but was also indirectly related to funding, as public perception of citizen science was reported to affect its funding. Similarly, participant retention, data continuity and university engagement were all bilaterally related to funding, project capacity, and support.

‘Knowledge sharing’ was not classified as an individual theme but was prominent across many other themes. For example, there were calls for better reporting and knowledge sharing to improve citizen science awareness, volunteer retention, confidence in the data, attitudes towards citizen science, and collaboration with educational institutions. In these instances, ‘knowledge sharing’ was spread across these other emergent themes.

Similarly, ‘open data’ appeared in five responses directly and four others indirectly. For example, under ‘participant acknowledgement’, responses noted that participants should have access to their own data to ensure the program is ethical and to improve retention. Related, a lack of confidence in data was the second most common theme and influenced other themes including attitudes towards citizen science, funding acquisition, technology, and volunteer retention.

The most prominent themes, including ‘attitudes’ and ‘data collection and confidence’ consistently arose in the responses to most survey questions (Fig. 3). Yet, responses to the question of challenges and barriers in citizen science disproportionately influenced emergent themes.

**Fig. 3:** Heatmap visualisation of the frequency of theme-related keywords in participant responses. The colour intensity and numerical values represent the prevalence of each thematic area, revealing which themes dominated specific questions and identifying patterns in participant discourse across the survey. The full list of keywords associated with each theme is provided in table S2 in the supplementary materials.

An overview of the themes, challenges and opportunities with illustrative quotes identified through the analysis, as well as how they align with the 10 Principles of Citizen Science (5), is included in table 1 below.

**Table 1:** Summary of themes, challenges and opportunities. Abbreviations: CS = Citizen science.

| Themes (number of responses) | Challenges | Opportunities | Illustrative quotes | Alignment with 10 Principles of Citizen Science |
| --- | --- | --- | --- | --- |
| **Recruitment and awareness (43)** | 1. Recruitment of new participants into CS projects.<br>2. Lack of public awareness of projects and how to become involved.<br>3. Lack of support and resources available for researchers recruiting volunteers.<br>4. Perceived inaccessibility to CS projects by non-scientists. | 1. Employ marketing and science communication strategies to increase awareness.<br>2. Target schools for recruitment.<br>3. Communicate benefits of participation and consider the needs of participants.<br>4. Providing tools to connect volunteers to projects.<br>5. Build relationships and collaborations for meaningful engagement. | “There is a lack of infrastructure to support making it easier for EMCRs and community to connect” [R15]<br>“People hear ‘science’ and think if they are not a scientist they can’t get involved.” [R34]<br>“Excellent support from marketing experts and teachers to help attract and retain participants really effectively” [R13] | P1 - active involvement. |
| **Data collection and confidence (38)** | 1. Widespread concerns of poor data quality or perception of poor data quality (28 responses).<br>2. Poor data continuity due to volunteer retention, funding, and unreliable long-term expert collaboration, for example university researchers on short-term contracts. | 1. Robust data validation, including standardised training, quality control, and data management.<br>2. Ensure data is shared according to FAIR principles (48).<br>3. Use standardised ways of reporting methods (including data collection).<br>4. Communicate data validation work to enhance trust and usability.<br>5. Potential for scalable data collection if challenges are addressed. | “Provision of open-access databases, QC protocols, and appropriate metadata standards can help with data validation and reliability.” [R11].<br>“There is a genuine lack of support for citizen science at both state and commonwealth levels which means it isn't considered a valid data source” [R40].<br>“There has to be a way to leverage the ability of citizen science to be almost everywhere at once to help inform research.” [R10]. | P1 - active involvement.<br>P2 - genuine outcome.<br>P6 - controlled limitations.<br>P10 - law and ethics. |
| **Project support and capacity (26)** | 1. Lack of centralised support for operationalising robust CS projects, for example project management positions at universities or not-for-profit organisations.<br>2. Lack of electronic and online infrastructure that is easily usable by citizen scientists. | 1. Provide centralised services to support project management, ethics and data management.<br>2. Develop institutional infrastructure that lowers the barriers for starting citizen science projects and supports project implementation and connection and engagement with communities.<br>3. Ensure the public are involved in all stages of the research, and there is transparent reporting about project capacity | “For some species-specific issues, there are no websites, databases or other resources available to promote and support data collection” [R14].<br>“A general hub with these [supportive] resources (national and international) would help find these resources” [R35].<br>“(Creating) enabling conditions for self-managed science” [R46]. | P4 - multi-stage participation.<br>P6 - controlled limitations.<br>P10 - law and ethics. |
| **Attitudes (25)** | 1. Perception of science as boring hinders citizen engagement.<br>2. Perception of CS as not robust.<br>3. Universities can become disinterested because of a lack of direct funding for CS activity.<br>4. Scientists and medical professionals may think CS is 'outreach', not research. | 1. Increased and improved outreach and information informing citizens about aims and opportunities to be involved.<br>2. Government is interested in CS because it's seen as inexpensive.<br>3. Create more consistent and clearer language to describe CS across disciplines.<br>4. Improve people's attitudes by conducting research on community attitudes toward CS; improving quality of projects, making benefits of CS clearer; improving information dissemination; and improving<br>industry alignment. | "Less authors lead to a more prestigious paper and higher recognition" [R43].<br>[In Australia] "in health and medical research, we talk more about 'consumer engagement, consumer involvement and consumer led' rather than citizen science. So there are also language issues to contend with" [R15].<br>"Unless you work in science, most people have a feeling about science as something hard and boring and for geeks" [R15]. | P2 - genuine outcome.<br>P3 - benefits science & society.<br>P8 - participants acknowledged.<br>P9 - outcomes acknowledged. |
| **Involvement and retention (24)** | 1. Retention of existing citizen scientists for involvement in ongoing data collection.<br>2. Delay between data collection and research outcomes hinders sustained involvement.<br>3. Lack of funding and resources for ongoing engagement.<br>4. Reduction in data quality with loss of CS participants. | 1. Design projects that are well-organised to facilitate participation.<br>2. Employ systems and personnel to manage volunteers.<br>3. Identify, provide and communicate benefits to participants.<br>4. Provide agency to citizen scientists by giving ownership beyond data collection, and by collaborating with key stakeholder groups. | “When people are just getting into science they need a fairly fast turnaround time between data collection, analysis, and results. The long arc of a research project is difficult to stay engaged with when it’s just casual engagement.” [R39]<br>“I think active participation is also important: moving beyond data collection to engage community members in multiple steps of the research” [R1] | P3 - benefits science & society.<br>P4 - multi-stage participation.<br>P5 - feedback provided. |
|  |  |  | “Ensuring these [data collection] platforms are accessible to all skill levels and intuitive to use, would trigger ongoing participation of the public.” [R43] |  |
| **Individual capability and training (23)** | 1. Researchers lack training to involve citizen scientists, identify appropriate collaborating organisations, and design projects.<br>2. Training of citizen scientists can be insufficient to meet program goals.<br>3. Meeting training needs requires expanded funding models and | 1. University-run training for researchers on developing successful citizen science programs.<br>2. Community-driven training of researchers from experienced practitioners.<br>3. Formalised training standards for citizen scientists, including micro-credentials for complex topics which can have career benefits for | “Academics sometimes lack the time and the skills to engage with citizen scientists” [R21]<br>“Researchers not knowing how to find citizen science groups, ... engage with sometimes large groups of very diverse people, ... and how to ensure integrity of data and analysis in a way that will stand up to the rigorous peer-review process.” [R38] | P2 - genuine outcome.<br>P3 - benefits science & society.<br>P6 - controlled limitations. |
|  | supports.<br>4. Researchers lack the time for high quality CS engagement. | volunteers.<br>4. Universities create capacity and recognise the time required to work with the public to create and sustain citizen science projects. | “Universities could provide resources and training on stakeholder engagement or establishing collaborations.” [R21]<br>“Appropriate training can enhance reliability and transparency of the data, as well as improve job prospects for participants.” [R11] |  |
| **Funding (23)** | 1. General lack of direct funding for CS as a methodology across all fields.<br>2. Limits research partnerships: short-term grants cause problems for recruitment, training, and institutional development needed for | 1. Develop mutually beneficial, ethical and transparent partnerships with industry.<br>2. Include dedicated funding for CS in existing grant schemes, including grants that better fit the design of CS compared to traditional research projects. | “They put in a lot of effort and learn a lot, but then funding is done and people move on” [R39].<br>“Invasive species, loss of biodiversity, climate change, [and] bushfire risk do not have an end-date [like funding cycles do]” [R45].<br>“Harnessing the unique selling points of | P1 - active involvement.<br>P2 - genuine outcome.<br>P3 - benefits science & society.<br>P9 - outcomes acknowledged. |
|  | engaging volunteers in long-term CS projects.<br>3. Affects data continuity. |  | citizen-science (...) to attract more funding" [R20]. |  |
| **Inclusion and access (20)** | 1. People involved in CS can be seen as simply data collectors without authentic involvement in the research process.<br>2. Ethical data sharing and intellectual property rights, especially when working with Indigenous participants. | 1. Include citizen scientists in all stages of CS including identifying topics and priority setting to create better connections between the public and professional scientists.<br>2. CS can help the public be involved in science in a meaningful way, ensuring it aligns with the values and priorities of the public. | "I think one of the challenges is the 'power' structure of research, that doesn't easily lend itself to 'ideas/assistance from the uninitiated' (...) scientists are put on a podium for the work they do, and there is not really knowledge of how non-scientists might contribute. We need to make it possible for there to be a 'space for everyone' in science, and obvious, accessible pathways into all those spaces" | P1 - active involvement.<br>P4 - multi-stage participation.<br>P7 - open data.<br>P10 - law and ethics. |
|  |  |  | [R15].<br>“Creating guidelines that emphasize ethical data sharing, intellectual property rights, and co-design of research projects with Indigenous communities would promote more inclusive and respectful collaborations, ultimately benefiting both science and society” [R27]. |  |
| **Recognition of volunteers (20)** | Volunteers are under-acknowledged and require additional support or recognition to ensure they feel their contributions are valued.<br>Volunteers may not have access to the data they’ve contributed | Allocating resources to provide support and recognition can encourage ongoing involvement.<br>Rewards, reimbursements and co-authorship can provide recognition.<br>Better support for intermediaries between citizen scientists and | “Citizen scientists should be rewarded for their participation via certificates, newsletters, training” [R43].<br>“Volunteers need to have better access to the data they themselves collect, because it is THEIR data (normative reason), | P4 - multi-stage participation.<br>P5 - feedback provided.<br>P7 - open data.<br>P8 - participants acknowledged. |
|  | - implies an ethical issue of public data availability and impacts retention. | researchers to support volunteers, such as community organisations.<br>4. Citizen science volunteers can co-manage projects alongside researchers. | because they care about and for data and not having access is detrimental for volunteer retention (instrumental reason) and because granting access could allow volunteers to do more with data (such as education and advocacy outside of the citizen science project's scope).” [R16]<br>“I think a formal intermediary role which could work between the parties would be hugely beneficial. Again, this could involve different levels of service/support based on needs of the group, but the focus should always be on multi-directional value exchange (citizen scientists learning<br>and acknowledged contributions, scientists gaining additional data and insights, broader community receiving value from new knowledge)." [R38] | P10 - law and ethics. |
| **Technology and innovation (16)** | 1. Redundant apps without innovation: many similar apps for CS can cause participants to be overwhelmed or bored.<br>2. Novel apps are technically difficult and expensive to build, with little support available. Difficult to keep up to date (software, security and interoperability). | 1. Technology has a crucial role in enhancing data accuracy, scale, participation and accessibility, ranging from visually impaired people to remote communities.<br>2. Support from corporations or governments in digital innovation can improve user experience and data interoperability amongst programs. | "Participants say "not another app" - [CS should] find innovative approaches that increase diverse participation, retention and enjoyment for all involved" [R29].<br>"There are already apps for almost everything you can think of nowadays, but nowhere to find an updated catalogue." [R7].<br>"Once [citizen science] is accepted more we can rapidly expand the scope of work | P2 - genuine outcome.<br>P3 - benefits science & society. |
|  | 3. It is difficult for some to find relevant CS apps based on their interests, despite there being many available. | 3. Central hubs that provide resources to help build digital infrastructure for CS projects and for participants to find CS projects. | and possibility of what can be achieved, especially with the use of AI [Artificial Intelligence].” [R20]<br>“More engaging, interactive mediums like apps that have dedicated teams updating and refining the front-end to ensure users can utilise the tools successfully.” [R7]. |  |
| **Educational institutions (15)** | 1. Lack of awareness of the benefits of CS to school students and staff.<br>2. Difficulty maintaining continued student involvement in projects. | 1. CS is an important tool for engaging and motivating students in science and solving real-world problems.<br>2. Strong partnerships with universities, including opportunities to build CS into curriculum and award | "School-based citizen science programs are a win-win (...) [schools] benefit from engaging with authentic scientific practices..." [R39].<br>"Appropriate integration of education [is important] (...) citizen scientists should be | P1 - active involvement.<br>P3 - benefits science & society. |
|  |  | certifications. | rewarded for their participation via certificates, newsletters, training..." [R43]. |  |

### Illustrative case studies

Survey participants shared 45 citizen science initiatives and an additional four were identified by co-authors. Seventeen STARDIT reports were created from the 49 examples, which were used to help select the final nine illustrative case studies (Table 2). Of the nine case studies, we invited people from all the initiatives to validate the data about them in the STARDIT reports, with 7 not replying, and 2 validating the data.

**Table 2:**
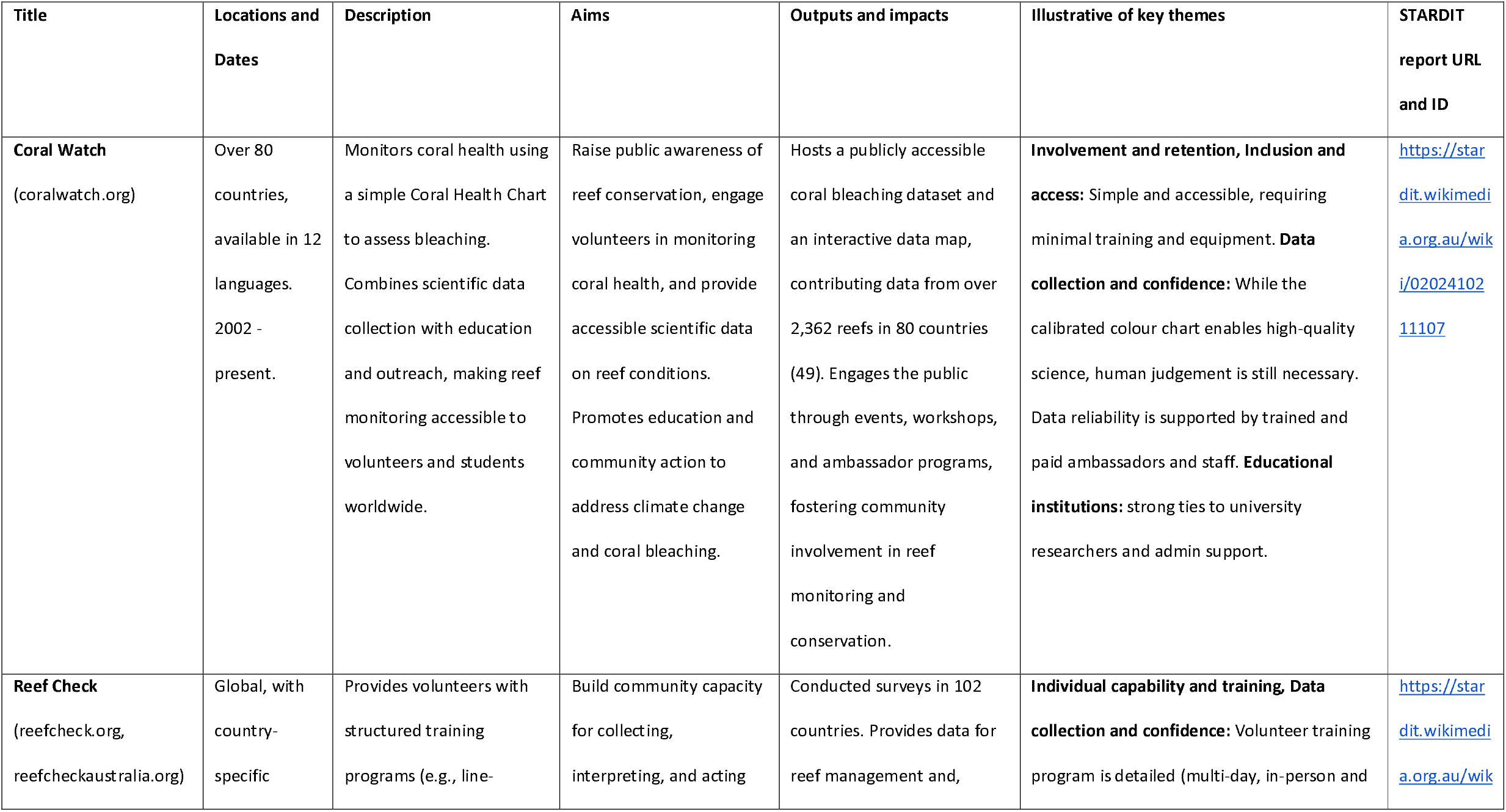
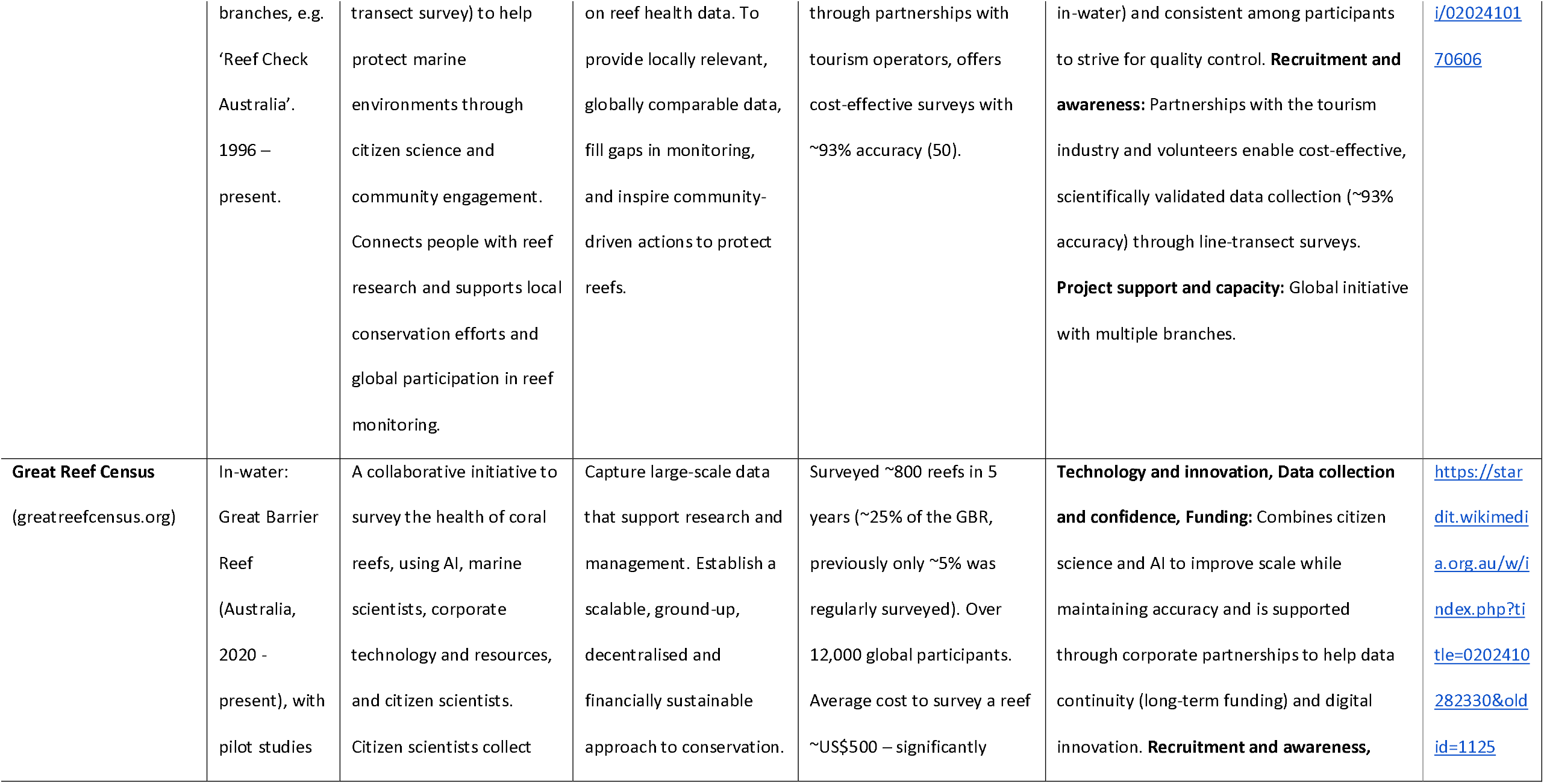
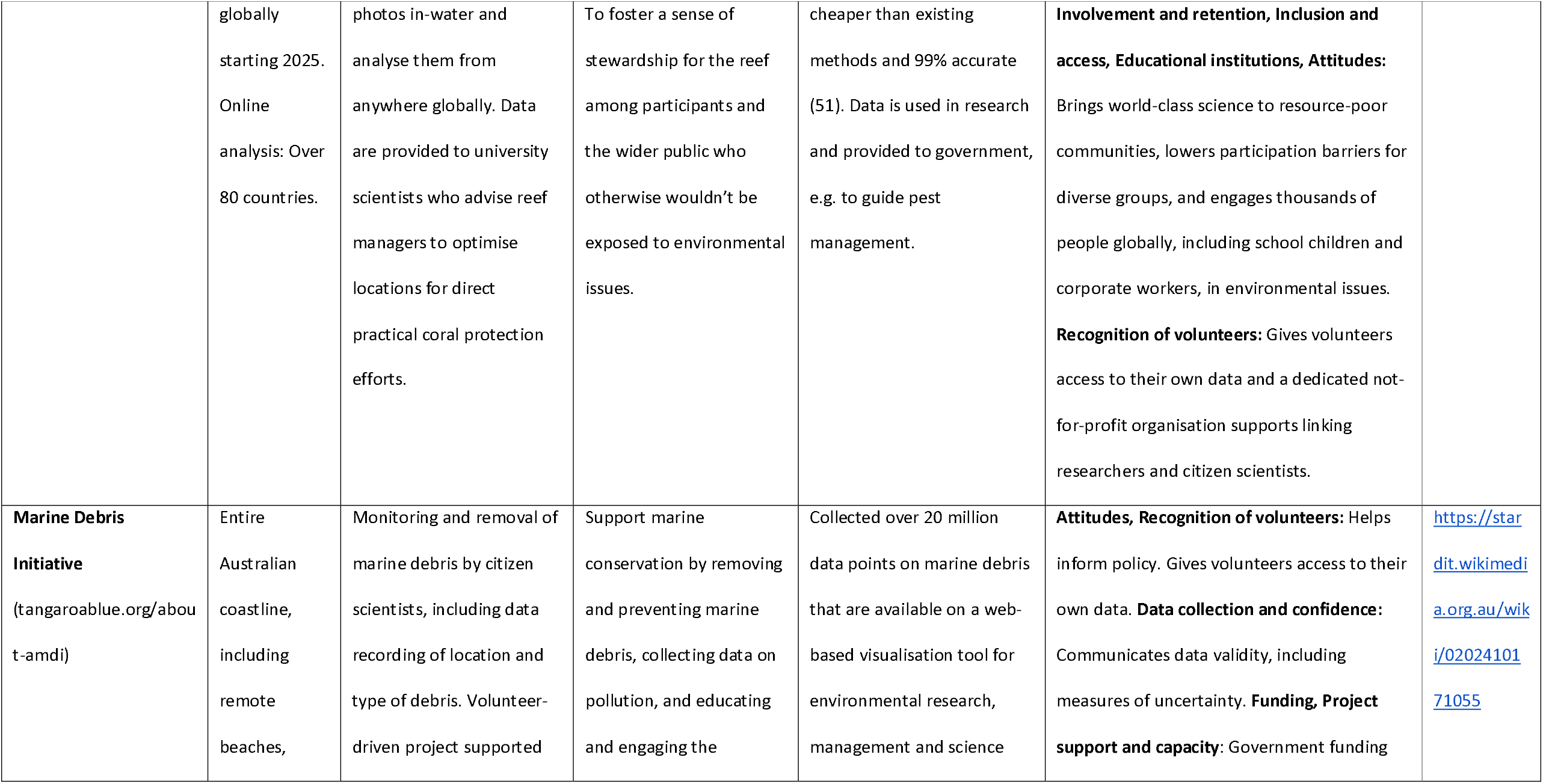
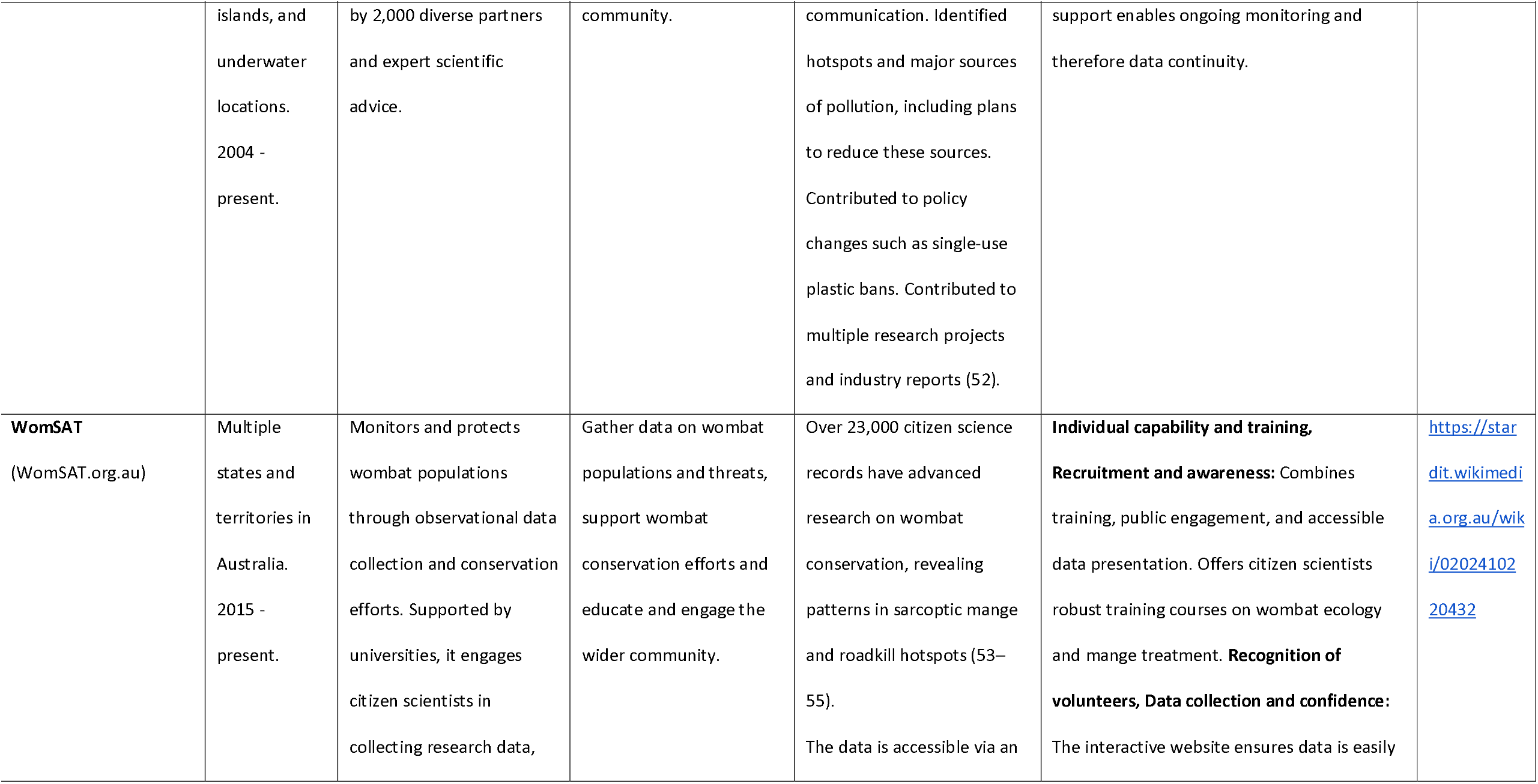
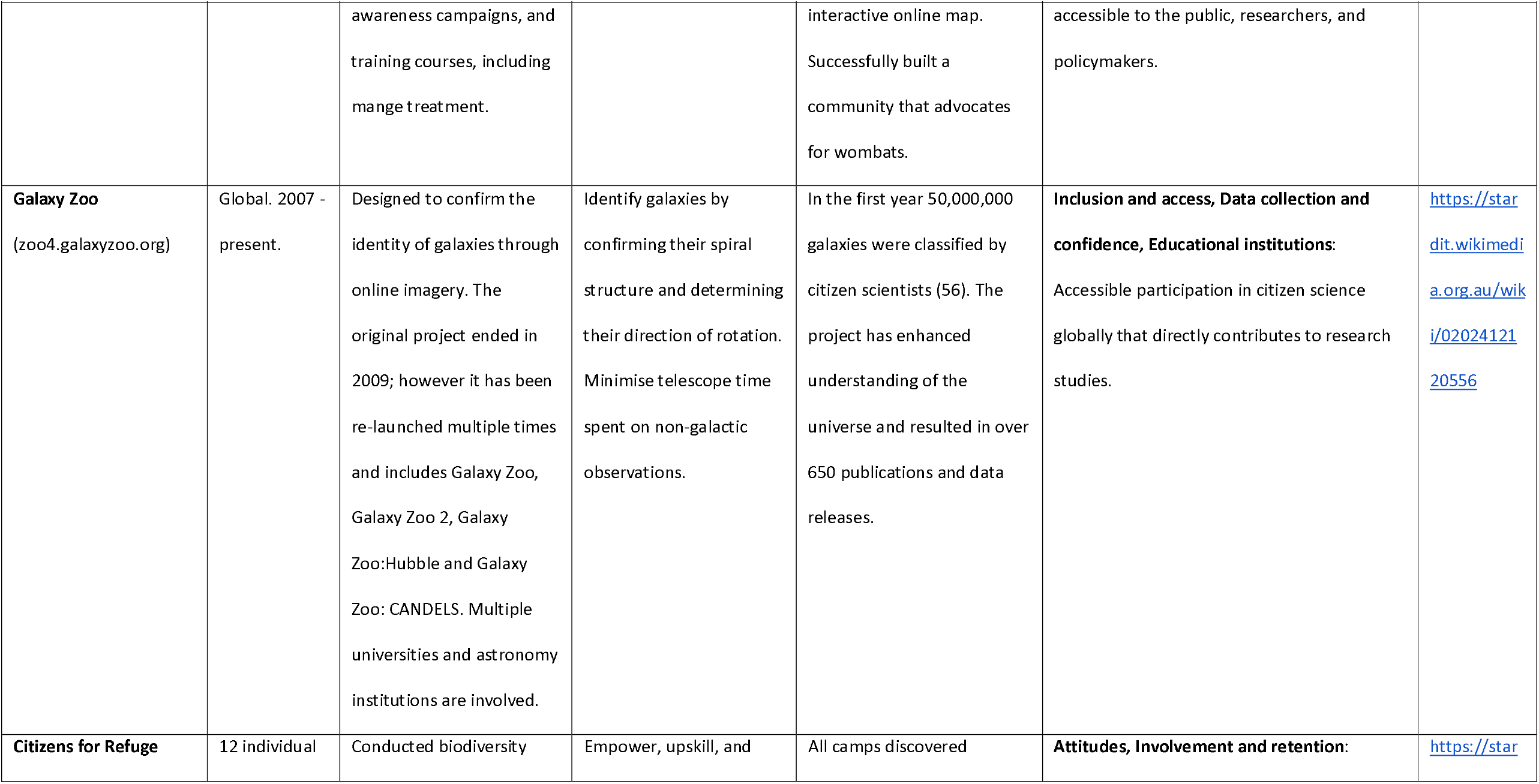
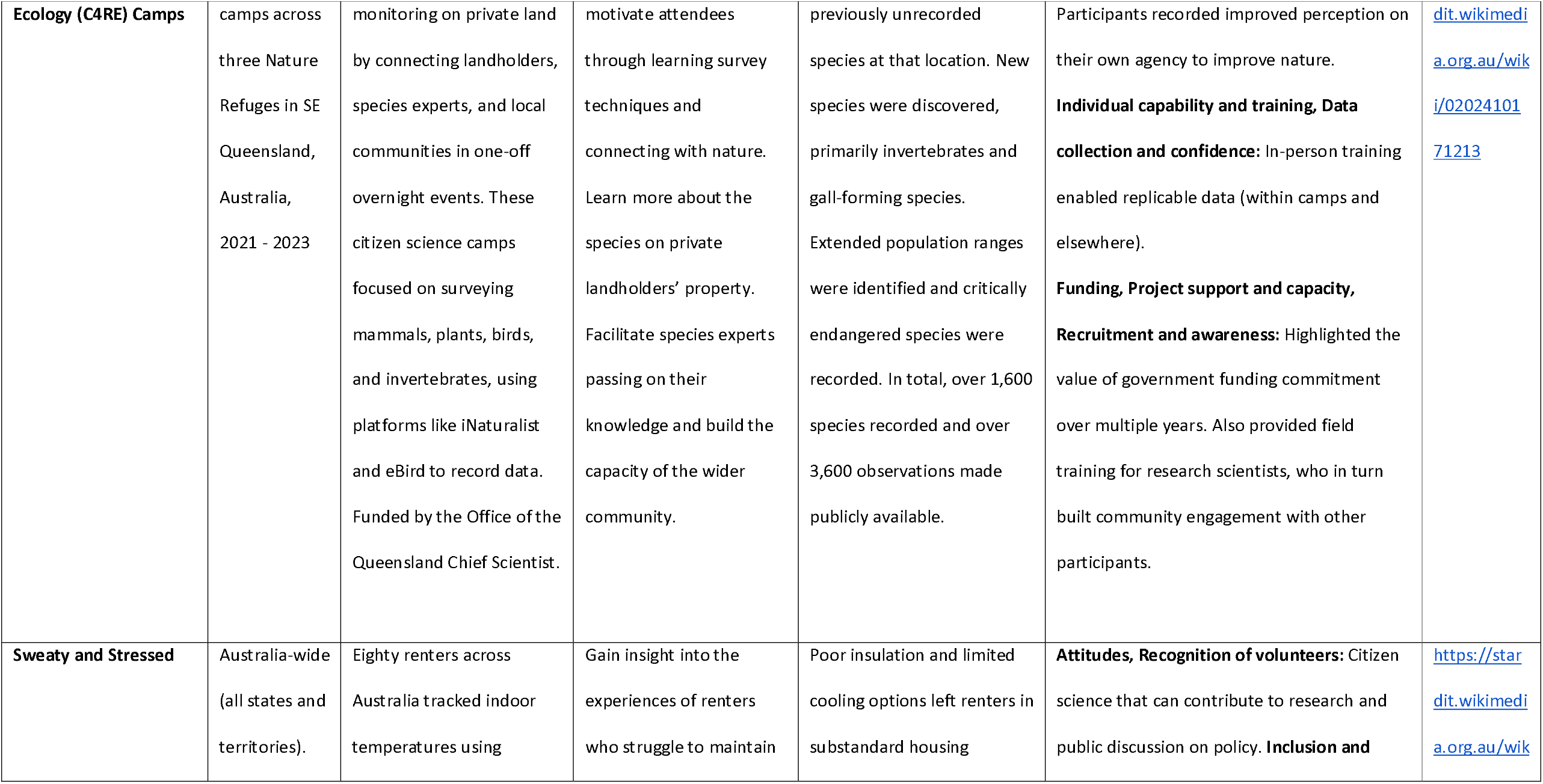
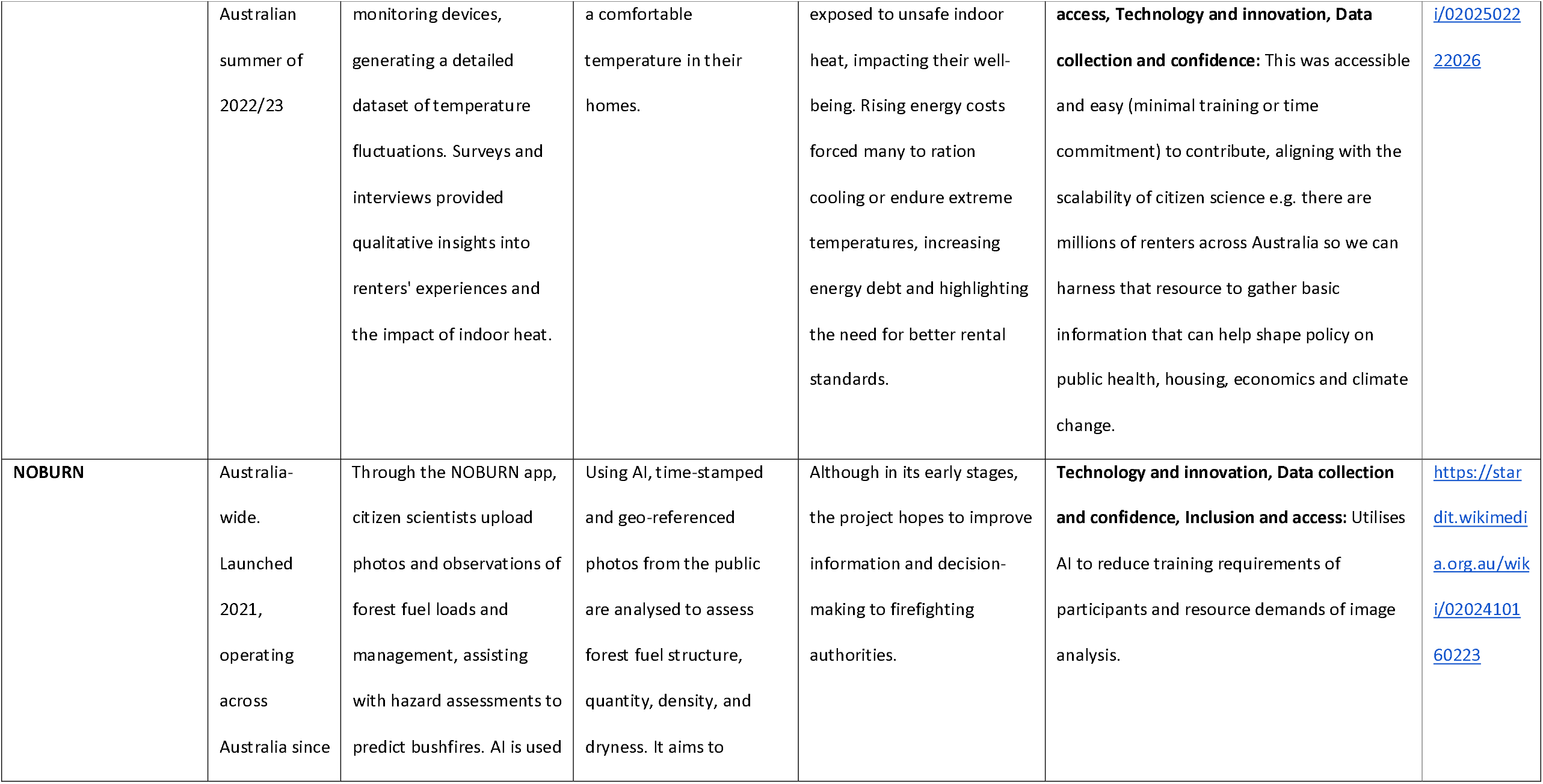
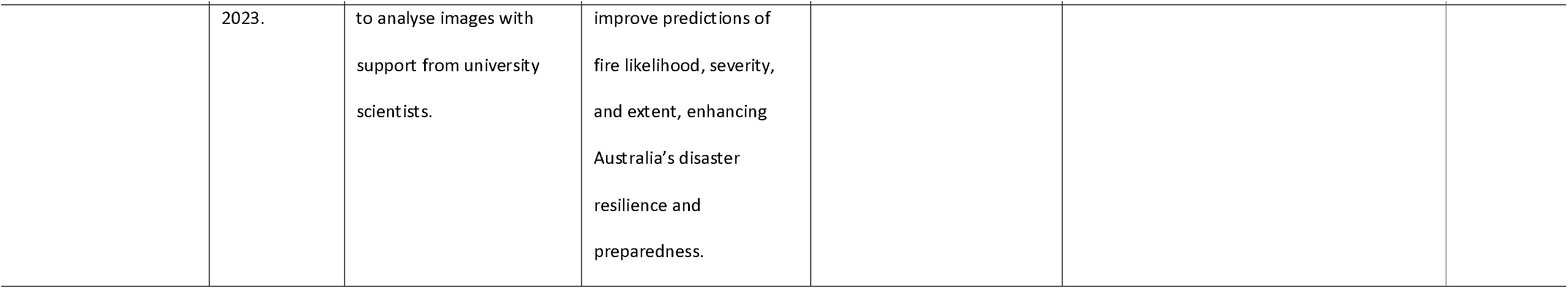
Illustrative case studies.

A cross-case analysis of the nine diverse citizen science projects from the illustrative case studies highlights key patterns across citizen science initiatives and provides examples of technologically integrated and collaborative approaches to enhance sustainability, data quality and public involvement. Projects varied in accessibility and citizen scientist commitment. For example, three projects focused on coral reefs but took diverse approaches. CoralWatch enables broad, low-barrier participation to in-water monitoring; Reef Check requires extensive training to support data reliability; while the Great Reef Census enables involvement in reef conservation from anywhere globally.

The integration of AI, as demonstrated in the Great Reef Census and NOBURN, reduced reliance on expert training, and enabled broader involvement. The use of AI to analyse photos uploaded by citizen scientists also reduced manual data processing, minimising human error and improving data accuracy.

Many initiatives foster deeper community stewardship by emphasising individual involvement and education, such as WomSAT and Citizens for Refuge Ecology (C4RE) Camps. CoralWatch, WomSAT and the Marine Debris Initiative are examples of how interactive online platforms can be used to involve and educate volunteers and give them access to data they have collected. These platforms are presented in user-friendly, and accessible formats, such as interactive maps.

Collaboration between paid scientists, unpaid volunteers, and partner organisations is highlighted as a key factor in project sustainability. However, funding models can differ. Corporate partnerships helped support the longevity and digital capabilities of the Great Reef Census, whereas C4RE Camps and other projects showcase the immediate benefit of short-term government grants. These variations highlight the evolving nature of citizen science, where balancing accessibility, data reliability, technological innovation, and funding stability remains a central challenge for long-term impact.

### Participation experience data

We collected data about people’s experience of being involved in this project, with 55% (15/27) responding ‘I felt I could be involved in all tasks of the project’, 45% stating ‘I felt I could be involved in some tasks’ and no one responding ‘I felt there was no opportunity to be involved in any tasks of the project’. When asked ‘Would you recommend using STARDIT to report citizen science or other initiatives?’, 10 people responded to the question, and 90% (9/10) responded yes, with one ‘maybe’.

## Discussion

Our co-produced results identified key challenges for citizen science, including participant recruitment and retention, ensuring data quality and trust, short-term funding limitations, and lack of payment or formal recognition for contributors. Effective strategies included transparent data practices aligned with FAIR principles, inclusive involvement methods, sustainable funding models, centralised support infrastructure, and expanded training and career pathways for participants. Case studies also highlighted the potential of technology and innovation, particularly data platforms and AI, to enhance scalability. Here, we provide 10 actionable recommendations in response to emergent themes that project operators, funders, advocators and policymakers can enact to improve citizen science and increase its scope for benefit to society and the environment. We also place our key themes and recommendations in the context of broader financial, ethical, political and legal issues.

### Recommendations

1. **Strengthen institutional and government support, and develop publicly led ‘hubs’**: Developing infrastructure, funding mechanisms, and policy alignment is important for sustaining long-term citizen science programs. These may be achieved if governments provide long-term funding for ‘super hubs’ that support institutions to host multiple citizen science projects. These ‘super hubs’ would use centralised funding and infrastructure for administration, ethics, project management, data hosting and sharing, communications, reporting and evaluation. For example, a ‘super hub’ at a university could build partnerships with schools, communities, specialist clubs (such as dive clubs and natural history societies) and beyond, while sharing control of the individual citizen science projects with the public.
2. **Improve partnerships with schools and educational institutions:** Strengthening partnerships between schools and higher education institutes can increase the number of people involved and the exchange of skills and knowledge among students, teachers and researchers. This could be achieved by integrating citizen science into curricula and providing appropriate funding and resourcing, such as training for early-to mid-career researchers and education professionals on developing partnerships.
3. **Improve communication about citizen science:** Evidence-informed communication methods should be used to inform the public and raise awareness about citizen science opportunities and their benefits. Increasing research on people’s knowledge and attitudes toward citizen science will be important to identify gaps, involve more people and work in ways that align with their needs and preferences.
4. **Improve citizen scientist support and recognition**: Multi-platform tools, training, and project management should be used to support inclusive ways of working and ensure data quality. Improved mechanisms to recognise citizen scientists is important and may include providing remuneration, clearer guidelines on acknowledgement and authorship in research articles, and better reporting on the contributions of citizen scientists using tools such as STARDIT (45).
5. **Improve transparency about methodology and processes**: Developing standardised frameworks for citizen science projects is key, including planning, reporting and evaluation. Quality assurance processes should be linked to reporting standards. Information from these frameworks can improve data validation, quality control, and communication about data reliability. Case studies using such frameworks can help demonstrate the value and effectiveness of citizen science methods, and any associated participatory methods. Data collected or analysed through citizen science also needs publicly accessible validation of its accuracy and/or reliability, as a perception of low-quality data can hinder data usage, future funding, and participant retention. Project design should include funding for scientific expertise to address concerns about data quality, help develop robust protocols, and to validate data. This means involving scientists from the outset, rather than having organisations design citizen science projects in isolation. Improved transparency about how data was collected and analysed can help improve trust in citizen science and improve data reliability and usability.
6. **Follow FAIR data principles**: Ensuring that all data are findable, accessible, interoperable and reusable with clear reporting standards is critical for impactful citizen science (48). This requires building, hosting and maintaining digital infrastructure for collaboration, including working with trusted partner organisations such as the Wikimedia Foundation. Large open data aggregators for citizen science should be utilised wherever possible, such as the Atlas of Living Australia working with the Global Biodiversity Information Facility. All government-funded research should mandate FAIR data sharing. There were many learnings about data collection, sharing and analysis. For data collection and sharing, online tools for citizen science need to work on multiple platforms and be inclusive and accessible. While four of the case studies we presented shared data according to the FAIR principles, five did not. This aligns with findings from other research that warns against projects where people donate their ‘unpaid work’ for data that is then be privately owned (10,57). The use and promotion of large open data aggregators, such as the Atlas of Living Australia, can ensure that data can continue to be accessible and used even once the citizen science initiative itself is no longer sustained. Important considerations for these data aggregators include clear quality controls, centralisation of the database, and the ability to flag incorrect records and filter data.
7. **Improve public involvement in all stages of research:** The public should be involved in all stages of science, from topic selection to project implementation and decision-making. This involvement should be reported transparently. This can be achieved by implementing inclusive involvement strategies with sufficient resourcing, including funding and staff time. All grant funding for citizen science by the government should require demonstration of how the public can be involved at each stage of the project. To ensure citizen science is inclusive, grant funding should provide specific funding to pay people for their time for being involved and ensure taxation practices do not inhibit involvement.
8. **Expand training opportunities**: University-led and community-driven training, including formal credentials, could be offered to support both professional researchers and citizen scientists. Training and support networks should be established, including a community of practice for early-to mid-career researchers who are initiating, leading or working on citizen science projects.
9. **Encourage transparent and ethical research partnerships**: There is a need to improve ways for the public to be involved in setting priorities for both non-government and government funded research. Transparent collaborations between industry, government, and academia can improve the information available about how well stakeholders align with the values and principles of citizen science and ethical research conduct.
10. **Improve consistency and terminology:** Relevant governments and international organisations (such as the United Nations, World Health Organisation and the Citizen Science Global Partnership) should lead a process to ensure there is consistent terminology describing citizen science involvement and its underlying values and principles across all disciplines (e.g. health, environment and education).

### Considerations for broader societal context

#### Long-term funding required

Involving the public through citizen science is central to many formats of research, yet one respondent commented that “there is a lack of infrastructure to support making it easier for Early and Mid-Career Researchers (EMCRs) and community to connect”. Resourcing (including funding, staff time and infrastructure) from governments, universities and other stakeholders needs to be increased, and needs to be longer term. Such long-term resourcing can ensure that citizen science activities can be more impactful by providing sufficient central administration and data infrastructure, along with capacity building and training for both the wider public and people leading citizen science (for example, at educational institutes). Given uncertainty in global funding for science, aid and conservation, citizen science should also explore diverse funding sources - while maintaining transparency on competing and conflicting interests. The use of centralised hubs that support decentralised science may be an efficient mode of utilising novel long-term funding sources.

#### Funding for inclusive and efficient ways of working

Our results also indicate that while people are more willing to complete unpaid tasks in some areas of science (such as data collection), there are clear areas for which citizen science projects need to be directly funded and resourced. Responses to our survey suggest a hybrid model, where some people volunteer and others are paid to complete specialised tasks, such as project management and navigating ethics processes. Such paid tasks could be efficiently centralised by organisations, with expertise in project management and ethics shared amongst multiple citizen science projects. Responses highlighted that a lack of centralised support structure, including project management positions, were a challenge for operating successful citizen science. Importantly, while projects may plan to rely on volunteers, certain ways of working may exclude some people from projects where people are not remunerated. This may include those unable to afford to volunteer their time (including those with caring responsibilities) or those without access to certain technologies. Co-designing a project with multiple stakeholders can help ensure it is more inclusive, aligned with people’s expectations, and is realistic. Co-designing involvement plans in this way can help ensure adequate support is available for people to become involved. Important factors may involve budgets for payment and expenses, technical support, or if appropriate, emotional support.

We also acknowledge that our definition of ‘paid’ in this study is intentionally narrow and generally refers to either an hourly financial compensation or as part of a salaried position, such as administration staff employed by institutions. We note, there are other reward mechanisms such as honorariums and non-financial rewards that are more appropriate for some participants, some of which appeared in responses under the theme ‘recognition of volunteers’. Such non-financial compensation may include scientific library access, co-authorship, education, or conference attendance. People should be involved in deciding the kind of compensation or rewards that are most appropriate for them.

#### Legal and ethical guidance on the use of artificial intelligence is urgently required

Technologies, such as artificial intelligence, are making new ways of working possible as the cost, accessibility and usability of such technologies improves. Artificial intelligence will thus inevitably play an increasingly essential role in future citizen science by enlarging the scale of data that can be analysed, and assisting with improving the quality of data collection, analysis, dissemination and translation (58). However, the ethical use and environmental impact of such tools and technologies often remains unclear, and further legal and ethical guidance from both governments and other stakeholders is urgently required (59,60). Additionally, as with all methodologies, improved reporting on the use of artificial intelligence needs to be transparently shared, including the underlying code and training data.

#### Improved data about political and other influences on science is required

Politicisation of scientific findings is increasing, as is misinformation and disinformation that contribute to the ongoing “infodemic” (61). As citizen science increasingly contributes to large-scale data collection and public involvement in research, safeguards from the politicisation of citizen-led research are needed (62). Areas of research including climate science, public health, drug development, and mental health remain politicised. Improved transparency of data on the different stakeholders will aid public understanding of where policy is influenced by evidence, ideology or economics. For example, multiple political factors are negatively impacting citizen science initiatives on climate change, including the ability of citizens and institutions to collect, share and interpret essential climate data (63–67). Globally, there is increasing political and corporate interference in climate science. This has included the funding of misinformation via fossil fuel companies who also sponsor government policy (68). In some countries, such as the USA, China, and Australia, government policies restrict access to environmental information (69–71). The defunding of climate institutions, shifts in public communication under political influence (as seen in Australia) and the subsequent self-censorship of researchers all have a direct impact on citizen science (65,72,73). Globally, citizen science can continue to be a force to collect, share and inspire action, but the impact of our recommendations will be limited if the politicisation of scientific findings continues to proliferate.

#### Evidence-informed methods of research dissemination and translation are needed to uphold democracies

A key benefit of citizen science is the direct involvement of communities and the general public in science, providing awareness and direction for scientific research that reflects the interests of the public. Citizens within democracies should review and co-create the ‘enabling conditions’ for science, including clear definitions and distinctions of what is ‘dissemination’ and ‘translation’, and what is ‘protest’ (74). In 2022, the United Nations Secretary-General stated “climate activists are sometimes depicted as dangerous radicals. But, the truly dangerous radicals are the countries that are increasing the production of fossil fuels” (75). In this context, the scientific method must objectively catalogue the methods of disseminating and translating research (including protests), alongside cataloguing the vested interests of people working for industries and other stakeholders (such as lobbyists), and their influence on government policies. Tools such as STARDIT can be used to map such interests in a transparent and collaborative way (45).

For people to be involved in citizen science, they require certain ‘enabling conditions’, which include transparency and personal safety. Consideration should be given to the duty of care owed by organisations conducting citizen science, particularly in relation to the personal safety of participants. This includes safeguarding individuals from potential harassment that may arise during data collection, dissemination, or translation of research. Further clarity is required on where the duty of care is situated in citizen science, and who might share it, including legislative, executive, or judicial bodies. For example, lawmakers could either reframe protests relating to scientific data as a legal form of scientific research translation and dissemination, or as ‘illegal protest’. The risk of ‘strategic lawsuits against public participation’ (SLAPPS) is also significant, and may impact people’s willingness to initiate, participate in or disseminate and translate results from citizen science (76). Legislation is required world-wide to dissuade such methods.

By analysing various methods of research translation and dissemination (through proper reporting and evaluation), we can explore evidence-informed approaches for translating scientific knowledge into action to prevent crises such as irreversible climate change and mass-extinction, and better protect those involved in translating findings from citizen science. Better reporting of such data will strengthen our collective ability to learn about safe and successful methods of dissemination and translation.

### Study Limitations

The survey responses were analysed qualitatively because the sample size of 46 was not large enough to make useful statistical inferences. Most respondents were Australian, and although the results primarily reflect the Australian context and perspectives, the findings and learnings are considered broadly generalisable. There was also an unequal representation of fields. For example, ecology was heavily represented while some fields were underrepresented, such as health (despite its significant share of governmental budget spending), or absent, such as climate and weather projects. The survey likely targeted individuals already engaged in citizen science, which introduced a bias toward those who view it positively. Perspectives from stakeholders with competing interests who might be less supportive of citizen science may be underrepresented. While responses included a range of stakeholders, the sample was skewed toward academia. This overrepresentation may have narrowed the range of perspectives, which in turn may have affected emerging themes and potentially omitted key issues important to the public.

### Perceived gaps in the data

As part of the data analysis process, the co-authors discussed the results and agreed areas or themes where they had expected to see more data or responses. The co-authors expected ‘open data’ and ‘data sharing’ to be more prominent. Similarly notable areas where no examples of best-practice were shared were ‘weather’, ‘climate’ and ‘air pollution’, despite these being significant areas for citizen science globally (33). The environmental impacts of citizen science itself was a theme that was also absent, along with any recommendations for offset practices (77).

### Knowledge gaps and opportunities for further research

As part of this research, we have identified two key knowledge gaps and opportunities for further research. Firstly, while we suggest that many of the themes and challenges identified as part of this research are reflective of wider patterns across the world, there are also likely large differences across social, economic, technological, political and cultural contexts. Secondly, while the citizen-science experts surveyed as part of this research hold key insights into how the impacts of citizen-science can be improved, we recognize that other stakeholder groupings - including policymakers, people working for industry, and people from marginalised communities - hold important perspectives for strengthening their own involvement in citizen-science projects and improving the overall outcomes of citizen-science initiatives. Further research could explore both gaps by extending the approach applied in this research to new geographic contexts and stakeholder groups. A key element of this work could be to identify best-practices that could be scaled up, replicated and used to strengthen international collaboration to better leverage the transformative potential of citizen-science.

## Conclusion

Citizen science has the potential to play a key role in generating scientific knowledge and supporting environmental and social action necessary for addressing the complex and multifaceted facing the globe in the 21^st^ century. The importance of involving people in shaping the future of science in multiple disciplines (including health, environment and education) will only become more urgent as the climate heats and more ecosystems collapse, impacting the health and survival of multiple lifeforms. Providing opportunities, spaces and funding can enable researchers to develop experiences and build the long-term connections necessary to improve citizen science. Improving systematic documenting and reporting of methods, impacts and outcomes of involving people in science will enable a more evidence-informed approach to improving inclusivity and resilience in citizen science. Using such evidence-informed approaches can improve citizen science to benefit society, global health, and the very ecosystems on which all life depends.

## Acknowledgements

Acknowledged Supporters

Misty Neilson (Forever Wild Initiative; Burnett Catchment Care Association; Thriive)

Caren Queiroz Souza (Federal University of São Carlos)

Jennifer Loder (Great Barrier Reef Foundation)

Aslina Baharum (School of Computing and Artificial Intelligence, Faculty of Engineering and Technology, Sunway University, Malaysia.)

Janelle Bowden (AccessCR)

Alice Motion (School of Chemistry, The University of Sydney) Jodi Salmond (Reef Check Australia)

Karen Johnson and Diana Kleine (CoralWatch)

## Data about acknowledgements

All these acknowledgements can be found in the STARDIT report about this project, which can be found here: STARDIT.wikimedia.org.au/wiki/0202407220511.

## Data availability statement

In the interests of transparency (78), we have shared as much data from the surveys as ethically possible, allowing others to critically appraise our analysis, re-analyse the data, or combine it with other data. The full dataset is available on the Open Science Framework: https://osf.io/wbv9s. Additionally, all STARDIT reports created as part of this project are publicly accessible, and links to them are available in the STARDIT report associated with this initiative STARDIT.wikimedia.org.au/wiki/0202407220511.

## Supporting information

**Glossary**

**Table S1: Overview of characteristics of respondents**

**Table S2: The themes and keywords associated with each theme used for the thematic analysis heatmap**

